# Novel approaches to quantify defective interfering particle activity reveal a surrogate marker of interference in mammarenaviruses

**DOI:** 10.64898/2026.09.26.754696

**Authors:** Rebekah Honce, Inessa Manuelyan, Chloe Schiff, Emily Van Beek, Kanayo Ikeh, Allysen Henriksen, Philip Eisenhauer, Jason Botten

## Abstract

Defective interfering particles (DIPs) profoundly influence viral replication and host responses yet it remains difficult to quantify their activity routinely in the laboratory. Here, using lymphocytic choriomeningitis virus (LCMV) as a model mammarenavirus, we establish complementary approaches for measuring interfering activity of DIPs across a range of experimental scales. First, we optimized plaque-and focus-based interference assays that directly quantify DIP-mediated suppression of standard infectious virus replication through a modified neutralization assay. Second, in cell monolayers saturated with both DIPs and infectious particles, we identified a complete loss of expression of the viral glycoprotein and matrix protein, which are each encoded in the positive-sense orientation of the mammarenavirus genome. Notably, this loss of protein expression mirrors the inhibition of plaque formation traditionally seen in this high DIP setting. The observed DIP-mediated loss of glycoprotein expression is a surrogate marker of interference and was used to develop an alternative assay of DIP activity. This glycoprotein expression assay recapitulated interference detected in traditional high-titer plaque assays (i.e. loss of plaques when all cells are saturated with DIPs and infectious particles) while substantially increasing scalability. These approaches enable sensitive and scalable quantification of DIP activity for LCMV as well as Junín virus and expand our experimental toolkit for studying arenavirus population dynamics, pathogenesis and immunity, and may help advance DIP-based antiviral strategies. This study also provides mechanistic details on how mammarenavirus DIPs interfere with infectious virus propagation.

## Introduction

Defective interfering particles (DIPs) are a hallmark of RNA virus replication. Preferentially produced during high titer infections, these unique components of the viral population were first described in the mid-twentieth century in groundbreaking studies with influenza virus [1, 2] and vesicular stomatitis virus [3]. As their name implies, DIPs are capable of viral interference and can effectively “shut down” traditional infectious virus replication. Often attributed to the action of truncated viral genomes, or defective viral genomes (DVGs) [4, 5], this interference can alter pathogenesis, antiviral immunity, and disease outcomes. Since these early findings, countless advances have been made in understanding their production, regulation, and role during experimental and clinical infections [6–9], which had led to their exploitation as a potential next-generation antiviral in the form of therapeutic interfering particles [10–13].

Identification and description of DVGs through whole-genome sequencing data and advanced bioinformatic pipelines has increased our understanding of their form, yet this reliance on genetic and genomic data has left a missing link between this sequence data and biological function of DVGs and the related DIPs [14–19]. Use of modern single-cell RNA sequencing has illuminated associations between DVG expression and host transcriptomic responses [20, 21]; however, these approaches do not work for all RNA viruses and are inappropriate for those viral families where we have yet to identify a genomic basis of interference. A lack of more traditional and high-throughput assay-based methods to measure interfering activity within viral stocks or experimental and clinical samples limits our ability to link the presence of DIPs with a corresponding interfering activity that may drive biological effects on pathogenesis—in essence limiting the connection between genotype and phenotype.

Present knowledge remains limited regarding the genomic basis, if any, of defective interfering activity, the structure of putative DVGs, and the impact of interference on pathogenesis and disease outcomes for mammarenaviruses, including the prototypic lymphocytic choriomeningitis virus (LCMV) [22–26]. To enable better study of this oft-overlooked component of the viral population, here we present three methods for the quantification of interfering activity for mammarenaviruses. Using a modified neutralization assay, we show robust and reproducible plaque-and focus-based methods to determine the interfering activity of DIPs in viral stocks, cell culture supernatants, and animal tissue. We also present a novel method to simultaneously determine infectious titers and interfering activity through the discovery of loss of glycoprotein immunostaining during interference. By providing easy to implement methods for the determination of interfering activity, our goal is to enable the field to more routinely capture and report this data alongside infectious virus measurements, particularly at this time, when the presence of DIPs and DVGs has become more appreciated for having a role in influencing viral disease processes and outcomes.

## Results

### Modified neutralization assay for determination of mammarenavirus interfering activity

Previously, we reported a modified plaque interference assay for the determination of interfering activity for LCMV [27, 28]. This method relies on UV-inactivation of standard infectious virus (while leaving DI activity intact) in virus-containing unknown samples, which are then plated in serial dilutions against a known concentration of infectious virus (Figure 1A). Enumerated plaques are then plotted against the dilution factor to calculate the inhibitory concentration for a half-reduction (IC_50_) in plaque formation (Figure 1B-D). While this method can finely resolve the inhibition of plaques and back-calculation of interference activity per milliliter (Figure 1E), putatively due to interfering activity of DIPs in the viral sample, a more high-throughput method was needed. A complementary method to the plaque assay, the focus assay, utilizes immunostaining to quantify viral particles per milliliter. We adapted this approach to scale down the interference assay from a 6-to 24-well format down to a 96 well format (Figure 2A) [29]. Here, after co-incubation of UV-inactivated unknown samples with a standardized dose of infectious virus, we performed immunostaining for the LCMV nucleoprotein (NP), the field-standard target for LCMV titration by focus assay (Figure 2B) [28]. As before, enumerated foci were plotted against the sample dilution for determination of the inhibitory concentration for a half-reduction in focus formation (Figure 2C, D), and calculation of interference activity per milliliter (Figure 2E). Using the same samples in the plaque interfering assay yields highly similar results (Figure 1E).

**Figure 1.**
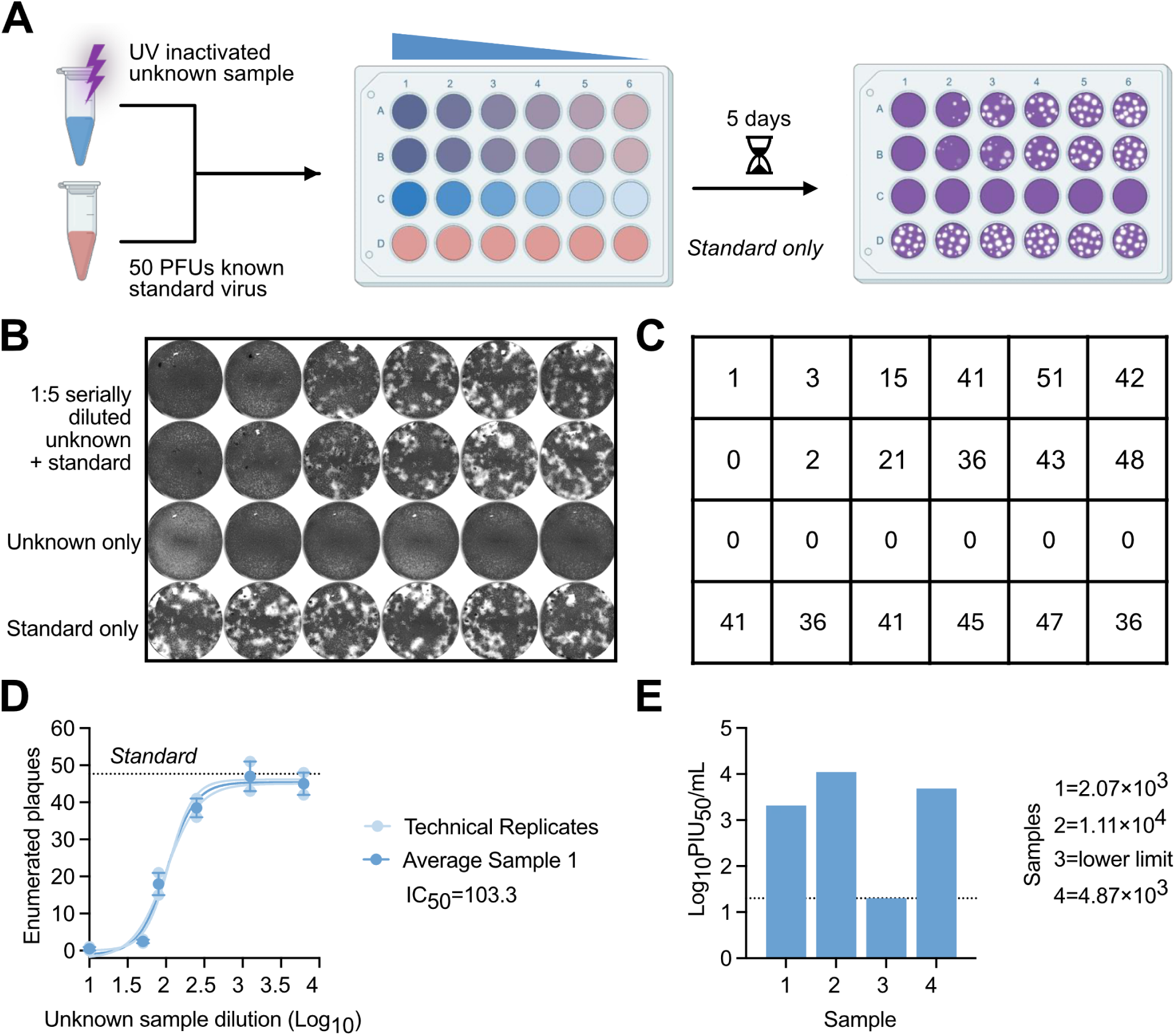
Traditional method of plaque interfering via modified neutralization assay. (A) Experimental schematic showing UV inactivation of an unknown sample and overlay with a standardized dose of infectious virus for determination of interfering activity via plaque interfering assay. (B) Representative crystal violet staining of plaque interfering assay test with (C) quantification. (D) Representative graph for inhibitory concentration-50 determination, showing each technical replicate and averaged sample used for calculations and (E) final expected results after dilution correction. Schematic in A designed with BioRender. Data in (D) graphed as individual technical replicates (light blue) or mean ± standard deviation and in (E) as absolute titer.

**Figure 2.**
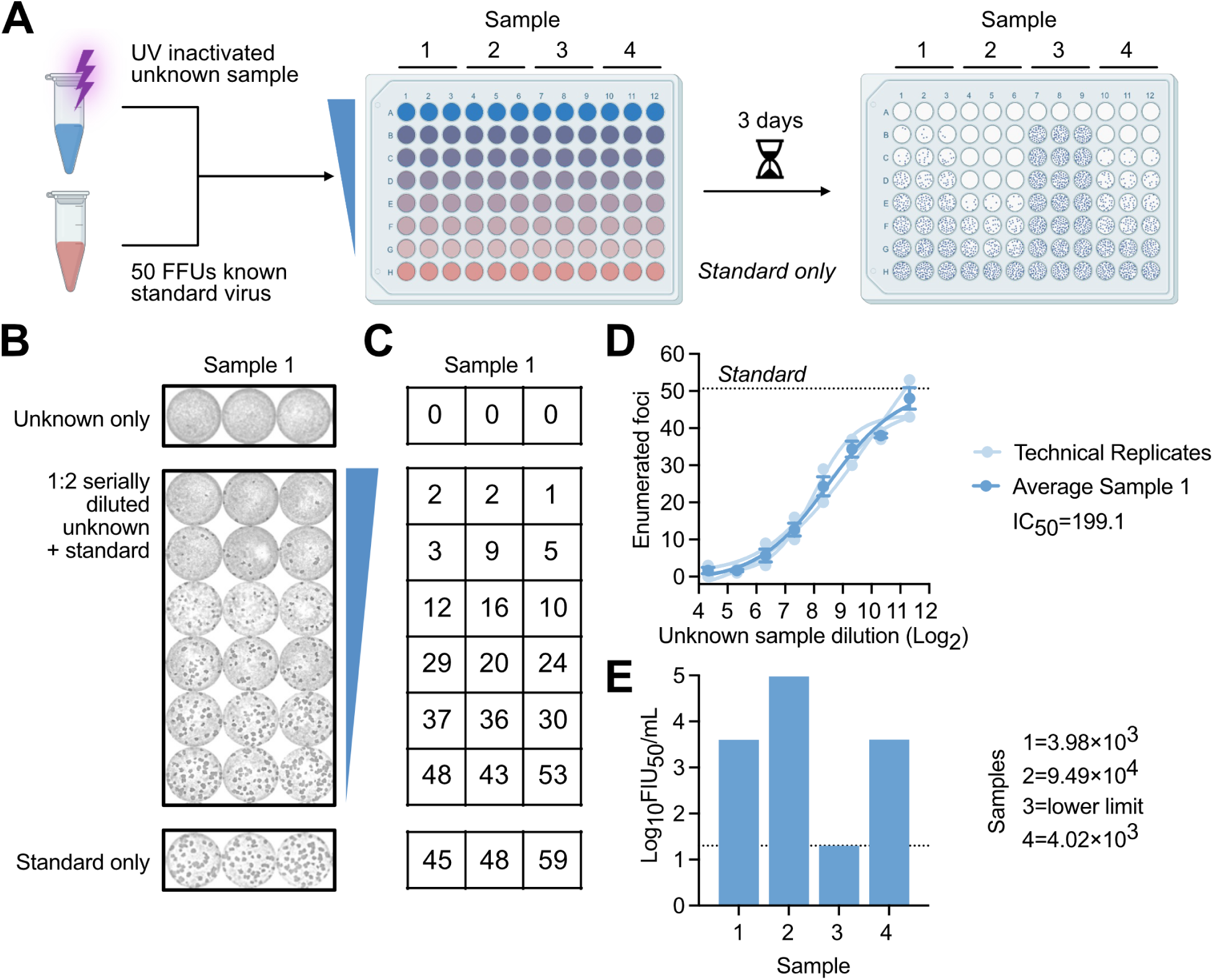
High-throughput method using miniaturized focus forming units. (A) Experimental schematic showing UV inactivation of an unknown sample and overlay with a standardized dose of infectious virus for determination of interfering activity via focus interfering assay. (B) Representative immunostaining for NP expression of focus interfering assay test with (C) quantification. (D) Representative graph for inhibitory concentration-50 determination, showing each technical replicate and averaged sample used for calculations and (E) final expected results after dilution correction. Schematic in (A) designed with BioRender. Data in (D) graphed as individual technical replicates (light blue) or mean ± standard deviation and in (E) as absolute titer.

### Loss of glycoprotein expression parallels interfering activity and enables simultaneous detection of interfering and infectious titers

While the plaque and focus interfering assay methods are highly sensitive and precise, they are time-and labor-intensive methods and unable to yield both infectious and interfering measurements in the same assay. This is further problematic for precious laboratory, clinical, or field samples when sample volume is limited. Thus, we sought to develop an assay that could simultaneously determine both infectious titer and interfering activity. We first observed distinct staining patterns for nucleoprotein (NP) when compared to either the viral glycoprotein (GP) or matrix protein (Z) in serially diluted LCMV supernatant samples (Figure 3A). This initial finding suggested GP and Z expression is lost in settings where cell monolayers are fully saturated with both DIPs and infectious particles. Viral L polymerase staining was not attempted due to a paucity of L-specific antibodies and the relatively low expression of L protein throughout infection due to its enzymatic nature and cellular toxicity [30, 31]. Thus, we performed serial dilutions of virus samples of unknown titers, while following traditional mammarenavirus focus assay guidelines, including overlaying the monolayers with methylcellulose following viral absorption [32], then stained the monolayers 3 days later with antibodies specific for NP or GP (Figure 3D). As expected, NP expression in Sample 1 of unknown titer mirrored sample dilution—with high levels of bold staining in wells with the most concentrated amount of virus, which dissipated with serial dilution to yield countable foci (Figure 3E, F). In contrast, GP staining of Sample 1 was absent at the highest concentrations of viral inoculum and only became apparent upon dilution— consistent with the expected effect of DIPs (Figure 3G, H). Despite this difference between NP-and GP-stained wells at higher concentrations of Sample 1, focus counts at the limiting dilution yielded equivalent infectious titers (Figure 3I). Thus, this pattern provides a surrogate measure for DIP levels: high viral inoculum concentrations exhibit strong interfering activity, which suppresses GP expression, enabling indirect quantification of interfering particles in parallel with infectious titer determination. Our findings also suggest that when a cell is infected with both DIPs and infectious particles, primary transcription and translation of NP, which is encoded in negative-sense orientation and transcribed from the genomic RNA, occurs while translation of GP and Z, which are encoded in positive-sense orientation and transcribed from antigenomic RNAs, does not occur.

**Figure 3.**
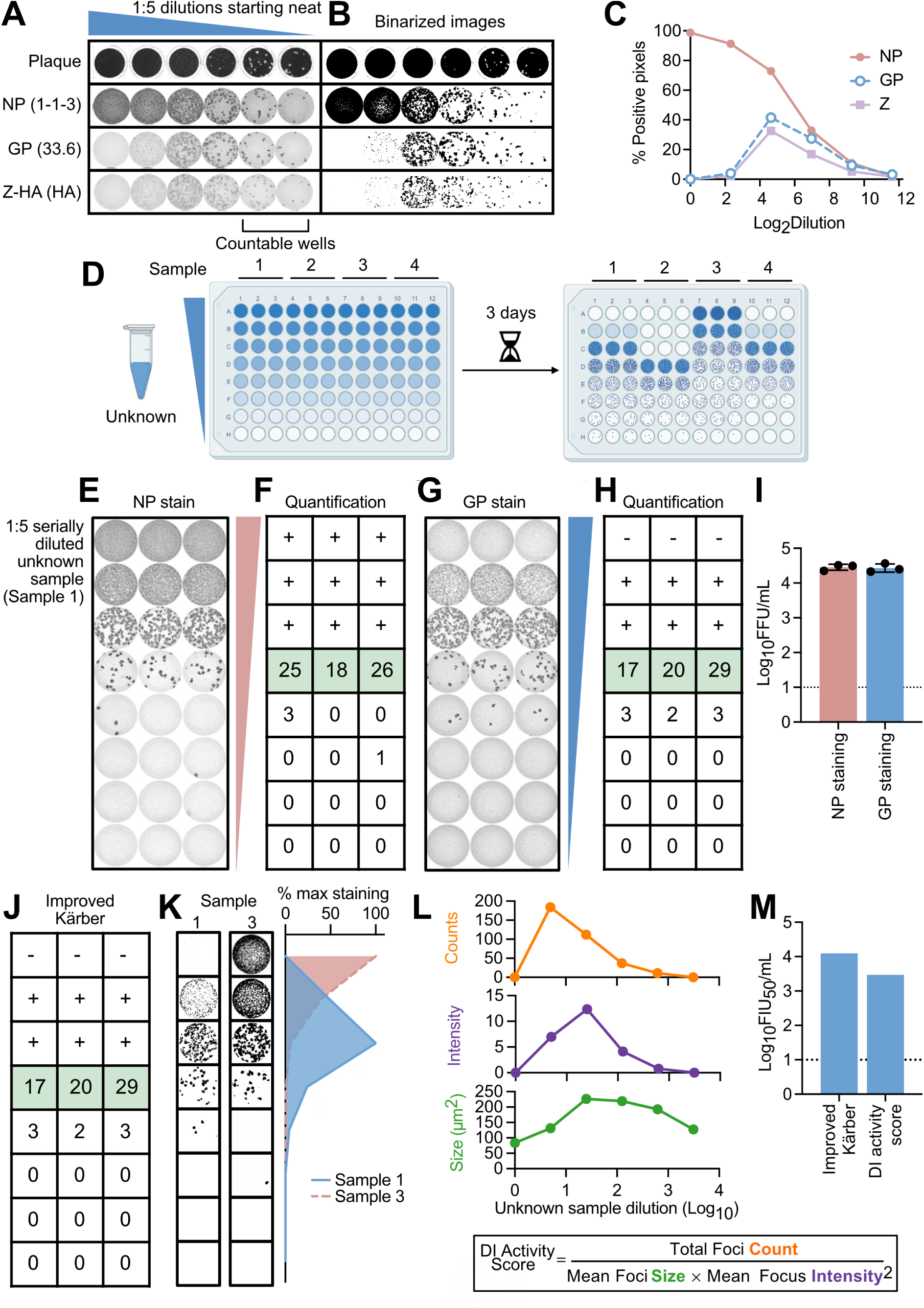
Dual determination of infectious titer and interfering activity via glycoprotein immunostaining. (A) Representative results of Vero E6 cells inoculated with 1:4 serial dilutions of rLMCV Z-HA and either overlaid with agarose containing media (plaque) then counterstained with crystal violet or immunostained with antibodies specific to the LCMV nucleoprotein (NP, 1-1-3 at 1:5000), glycoprotein (GP, 33.6 at 1:5000), or HA-tagged matrix protein (Z, HA 18181 at 1:2500). (B) Representative images in (A) were binarized and (C) positive pixels were graphed against sample dilutions. (D) Experimental schematic of focus assay; cells are first inoculated with serially diluted samples then immunostained with NP or GP antibodies (1-1-3 or 33.6, respectively) 3 days post-infection. (E-I) Representative focus assay results for (E) anti-NP staining and (F) quantification or (G) anti-GP staining and (H) quantification for Sample 1. (I) Results of calculated infectious titer of Sample 1 using first countable well highlighted in green on (F, H). (J-K) Representative calculation for interfering activity of Sample 1 using (J) the Spearman-Karber method aided by (K) maximal staining analysis performed in ImageJ. (L) Representations of raw data for total foci counted, total foci intensity, and mean foci size plotted across sample dilutions, which are the metrics underlying DI activity score calculation. (K) Comparison of two quantification methods for interfering activity determination. Schematic in (B) designed with BioRender. Data in (G, K) graphed as mean ± standard deviation with symbols representing technical replicates and in (I) as percent of black pixels in the region of interest.

We considered several ways to quantify the loss of GP staining as a measure of interfering activity. The most direct is via a visual inspection of tissue culture interference dose (TCID)-50 (Figure 3J, K) and we would recommend this method for relatively small numbers of samples. When a high-throughput option is necessary, a mathematical calculation incorporating the total number, size, and intensity of stained foci (Figure 3L) can be used to quantify DI activity scores. Similar to plaque and focus-interference assays, DI activity scores can be plotted against sample dilution to identify the dilution factor at which the DI activity score decreases by 50 percent, which is then standardized to a per milliliter value. Both quantification methods (Figure 3M) yield results consistent with those obtained using traditional plaque and focus-based interfering assays.

Instruments used to quantify stained foci report various metrics in addition to count such as focus size, intensity, or percent of well covered by foci. However, because no single metric can distinguish interference from total viral signal on its own (see Discussion) we developed a composite DI activity score. This mathematical approach accounts for the observable qualitative differences of focus size and staining intensity at higher versus lower virus concentrations, where increased DI activity generally results in smaller and fainter foci while reduced DI activity results in larger, more intensely stained foci (Figure 3G). To quantitatively capture these differences at each virus dilution, we based the DI activity score on three measurable properties of stained foci: total focus count, mean focus size, and mean focus intensity (Figure 3L). Focus count provides a measure of the number of detectable infection events, whereas focus size and staining intensity putatively correlate with the level of viral replication during infection events. Because larger and more intensely stained foci suggest more productive viral replication and potentially reduced DI activity, mean focus size and mean focus intensity were incorporated as denominator terms, with intensity weighted quadratically to increase the contribution of the large differences in staining intensity observed in wells exhibiting high- versus low-DI activity. Thus, the resulting DI activity score increases when a greater number of small, faint foci are present and decreases when foci become larger and more intensely stained.

In summary, our findings establish the loss of GP or Z staining as a surrogate marker of interfering activity that can be captured via focus assay. By changing the immunostaining target from NP to GP, this modified focus assay yields both an infectious titer and a measure of DI activity from the same wells, restoring access to interference information previously obtainable only through separate plaque-or focus-based interfering assays.

### Strengths of assays in varied experimental samples

We next applied these plaque-and focus-based methods to various experimental samples, including cellular supernatants from viral growth curves, serum samples from experimentally infected mice, and results of reverse genetics rescue of LCMV mutants. All samples were coded so to ensure research staff were blinded to sample identity until after analysis. First, we infected Vero E6 cells at either a low (0.0001) or high (0.01) multiplicity of infection (MOI) and collected supernatants at 24-, 48-, and 72-hours post-infection (Figure 4A). The infectious titer and interfering activity were determined by plaque assay and plaque interfering assay, respectively. As expected, we observed the predominant production of DIPs at high MOIs, and the loss of infectious virus production as interfering viral loads rise (Figure 4B-C). Next, we applied the focus interfering method for a high-throughput and sensitive assay for the determination of interfering viral load *in vivo* (Figure 4D). As the focus interference assay requires much less input material (50 µL) than the plaque interfering assay (250 µL), it is advantageous when sample volume is limited, such as mouse serum derived from a survival blood draw. In our application, we were able to see clear evidence of interference in serum collected from an individual, persistently infected C57BL6 mouse (Figure 4E), which can be graphed and quantified for between-group comparisons (Figure 4F, G). Finally, the GP staining method allows for fast screening of viral isolates that may be attenuated due to changes to DIP production compared to changes to other viral replication machinery (Figure 4H). Here, Samples 1 and 2 have similar infectious titers; however, they differ in interfering activity (Figure 4I). Conversely, Sample 3 has an approximate 1-log reduction in infectious titer, but no detectable interfering activity. Staining for NP expression would fail to show these differences and yield incomplete information on viral biology.

**Figure 4.**
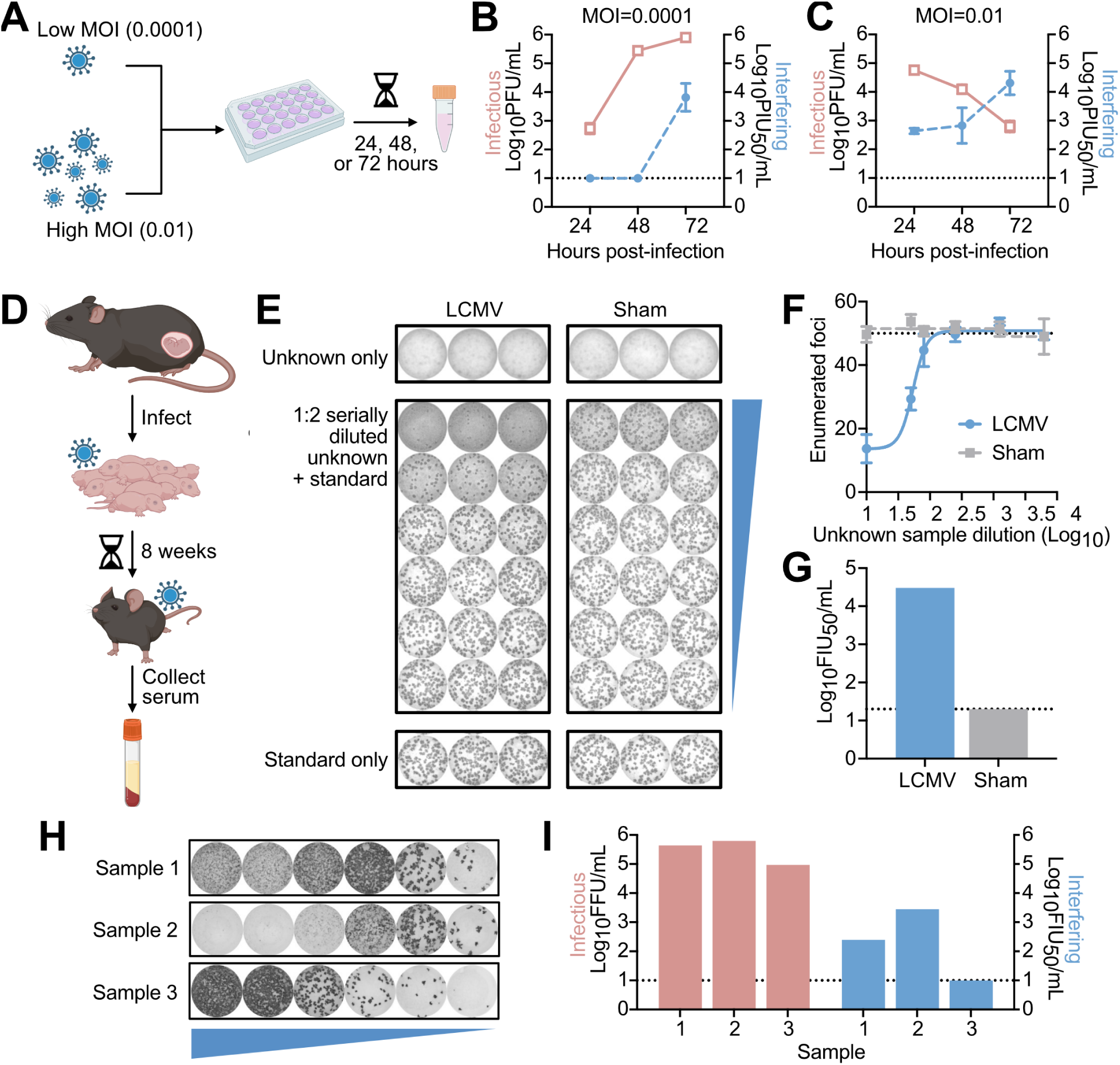
Application of foci-based method for experimental samples. (A) Experimental schematic showing infection of Vero E6 cells with low (0.0001) or high (0.01) multiplicity of infection (MOI). (B-C) Infectious titer and interfering activity results, determined via plaque assay or plaque interfering assay, respectively, of (B) low MOI infection versus (C) high MOI infection. (D) Experimental schematic for the generation of mice persistently infected with LCMV. (E) Representative staining of a focus interfering assay performed using serum harvested from an individual LCMV infected versus sham infected mouse. (F) Representative graph for inhibitory concentration-50 determination and (G) final expected results after dilution correction. (H) Representative results of 3 recombinant LCMV stocks (Samples 1-3) serially diluted in a focus assay and immunostained for GP expression. (I) Subsequently calculated infectious titers (red) and interfering actvity (blue) of Samples 1-3; interfering activity were determined using DI Activity Scoring. Schematics in (A, D) designed with BioRender. Data in (F) graphed as mean ± geometric standard deviation with symbols representing the geometric mean of 3 technical replicates and in (G, I) as absolute values. For ease of display, mouse experimental data in (D-G) is plotted for n=1 infected and sham infected mouse.

### Application of immunostaining approach across multiple viruses

Finally, we expanded our viruses of interest to include multiple strains of LCMV as well as the Candid #1 vaccine strain of Junín virus (JUNV) [28, 33] (Figure 5A-C). We observed that GP-mediated immunostaining can reveal strain and species-specific interference patterns. Notably, in contrast to the complete absence of GP signal in low-dilution wells of wild-type LCMV Armstrong 53b strain (Figure 5A; “Focus Assay – GP”), a recombinant LCMV incapable of producing DIPs [28] did not display reduced GP signal at these same low-dilution wells (Figure 5B; “Focus Assay – GP”) but resembled the NP immunostaining pattern (Figure 5B; “Focus Assay – NP”), supporting our observation that DIPs decrease GP signal. Repeating the focus assays with the JUNV Candid #1 vaccine strain (Figure 5C) shows a similar trend as wildtype LCMV Armstrong 53b, albeit with JUNV Candid#1 generating lower quantities of DIPs as evidenced by the right-shifted maximal staining for GP relative to NP for wild-type LCMV and JUNV Candid#1 [34]. Overall, when quantifying staining intensity for GP versus NP, the differences in apparent interfering activity may be inherent to these viral species or strains, or a result of growth conditions and timepoints (Figure 5D-F).

**Figure 5.**
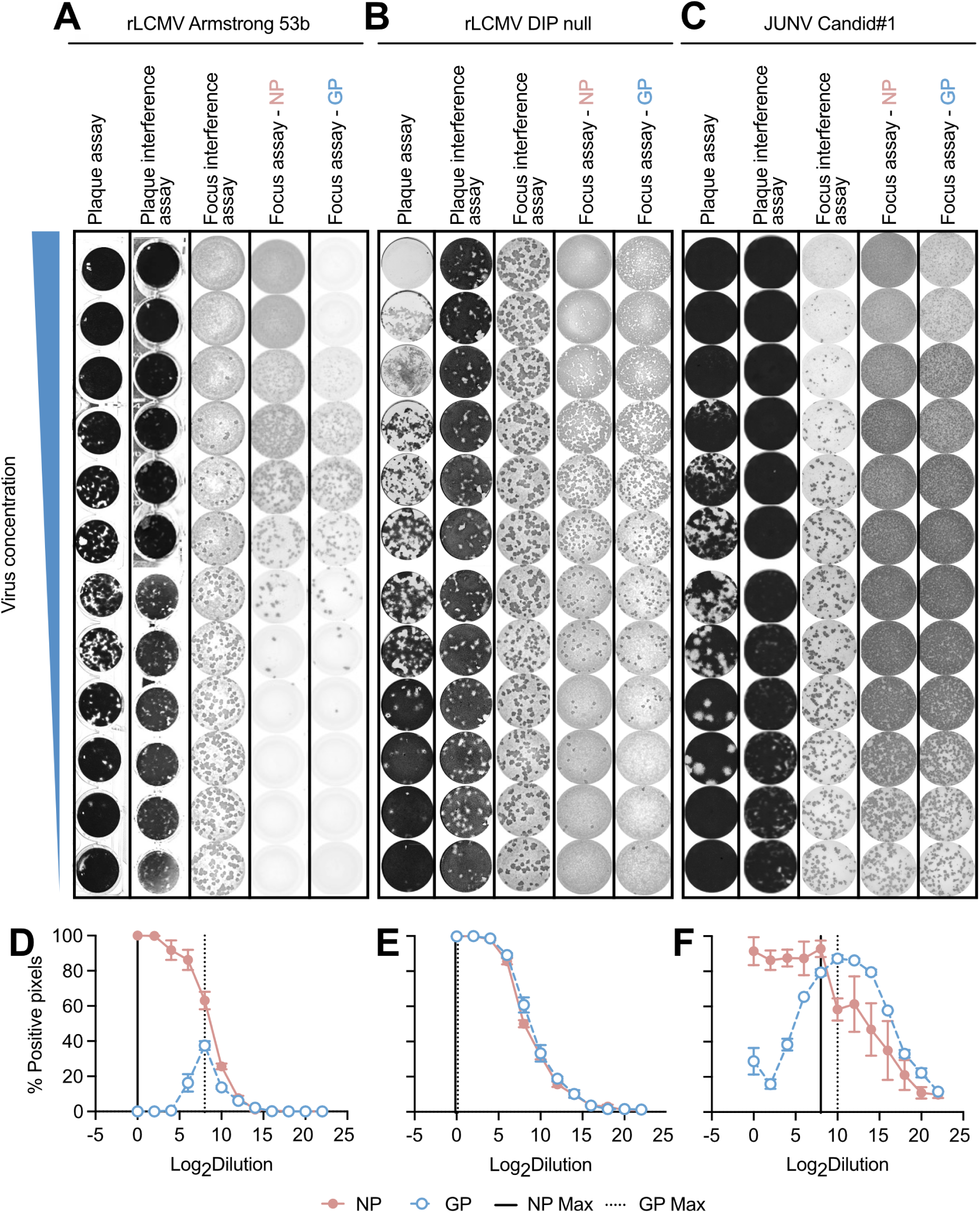
GP expression is restored in high-concentration sample wells of DIP null LCMV recombinant virus. Application of plaque assay, plaque interference assay, focus interference assay, and nucleoprotein (NP) and glycoprotein (GP) immunostain focus assays for (A) LCMV Armstrong 53b, (B) an rLCMV DIP null recombinant strain incapable of producing defective interfering particles, and (C) Junin virus Candid#1 vaccine strain. All assays began with neat sample then proceeded with 1:4 serial dilutions. (D-F) Quantification of foci staining intensity across (D) wild-type rLCMV Armstrong 53b, (E) rLCMV DIP null, and (F) Junin virus Candid#1 strain represented as a percent maximal staining per well and graphed as means ± standard error of n=2 or 3 technical replicates per virus. Experiment was repeated in triplicate with representative results of a single replicate displayed in (A-C). Plaque assay and plaque interference assay well images captured via an HP scanner, Cytation7, or CTL ImmunoSpot S6 Image Analyzer. Plaque assay and plaque interference assay were performed in 24-well plates, while the focus interference assay, and NP-and GP-immunostained focus assays were performed in 96-well plates. Equivalent volume per area of well was used to scale initial dilution to capture similar patterns of interference.

## Discussion

Though the importance of defective interfering particles in mammarenavirus biology cannot be overstated, there are a lack of high throughput, sensitive, and reliable assays to measure their activity. Here, we have modified our previously reported plaque interference assay to a focus interference assay amenable to high-throughput sample testing and developed a new focus assay using GP immunostaining which allows for the simultaneous assessment of infectious viral titers and DI activity. After testing these three assays and finding complementary results, we recommend the use of a single assay throughout an experiment, including the method for quantification. To determine the appropriate assay to use, the number and volume of available samples should be considered. When sample volume is not limited and there are a relatively small number of samples to be tested, the plaque interfering assay offers distinct advantages: it bypasses the need for immunostaining and specialized imaging equipment, relying instead on direct visualization of plaque morphology. Conversely, when plate scanning technology is available and sample volume is limited and/or there are many samples, the scaled-down focus interfering assay offers a fivefold reduction in sample input. And, in contrast to the plaque interfering assay, which requires one 24-well plate per sample with only a 6-point dilution series, the focus interfering assay can accommodate up to four samples per plate and supports up to a 10-point dilution series. This enhances both the resolution and throughput of viral quantification. Lastly, regardless of sample number and volume, transitioning from NP-based to GP-based immunostaining for the routine titering of LCMV will increase the yield of the standard focus assay. The GP-based focus assay enables both the precise quantification of infectious titer and reveals interference patterns indicative of the presence of DIPs which has historically been observable only in traditional plaque assays with high-titer stocks. While plaque assays inherently capture both infectivity and interference, their use has declined in favor of the more scalable focus assay. By adopting GP-targeted immunostaining, we restore the capacity to detect interference within a modern, high-throughput framework, enriching our understanding of viral dynamics without sacrificing rigor or increasing experimental costs and personnel effort. Then, if more sensitive quantification of interfering activity is required, follow-up with the plaque or focus interference assays allows for more reproducible and precise methods to formally quantify interference.

In lieu of a formal quantification of interference by a focus interference assay, the GP-based focus assay can be also used to quantify DI activity by a straightforward adaptation of a TCID_50_ measurement or by calculating a DI activity score for each well, then determining the dilution at which this score is reduced by 50 percent (Figure 3J-M). We developed this score using the foci metrics commonly reported by plate scanning instruments. The choice to combine foci metrics into a single score, rather than plotting any one directly as an indirect measure of DI activity, reflects the limitations of the individual measurements returned by automated foci analysis. Specifically, focus count alone reflects the total quantity of virus as much as its interfering activity, and at the highest sample concentrations where DI activity is greatest, dense or coalescent staining can be fragmented into numerous detected objects, such that the most concentrated well can also register a high rather than a low count. The percent of well area covered by staining is similarly ambiguous, as fields of small, lightly stained foci due to DI-mediated suppression and those with large, intensely stained foci can yield comparable well coverage; this metric therefore also fails to capture the effects of DI activity apparent by eye. A more intuitive approach would be to plot total signal intensity of foci per well against sample dilutions, where low signal intensity in the first few dilutions of sample would serve as an indirect measure of high DI activity. However, this approach typically yields a bell-shaped, biphasic distribution, as increasingly dilute samples that no longer contain high viral titers nor DI activity (as verified by traditional plaque or focus interference assays) also display low total intensity. In principle, the relevant IC_50_ can be calculated from the rising limb of such a curve, but the biphasic distribution is disadvantageous for two reasons. First, accurately fitting a biphasic curve with relatively few data points (in our example, 6 datapoints stemming from 6 dilutions) is difficult. Second, in experiments comprising many samples with diverse DI activity levels, biphasic and monotonic distributions will be intermixed, and unless the appropriate curve fit is considered for each sample, the resulting IC_50_ values may be inaccurate or nonsensical. In these cases, manual exclusion of irrelevant datapoints from the descending tail of biphasic curves would be required, which is feasible with a relatively small number of samples, but may be impractical for high-throughput experiments. By considering count in combination with focus size and intensity, we found that the composite score captures the qualitative distinctions between foci in wells affected versus unaffected by DI activity, while yielding a measure that decreases monotonically with dilution across most samples, reducing the manual curation required at scale. A useful feature of the DI activity score is that samples with very high interfering activity, which yield no quantifiable foci at the lowest dilutions, return no score in those wells and are thus excluded from the curve fit automatically without the need for manual intervention. An important limitation is that a subset of these samples with high DI interference may still exhibit a rising limb, due to the detection of very low numbers of foci. The composite DI activity score therefore greatly reduces, but does not eliminate, the manual exclusion of leading points before curve fitting for some samples.

The reduced GP expression pattern we observed in low-dilution, high-virus concentration sample wells was also apparent with Z matrix protein immunostaining (Figure 3A). However, the limited availability of Z-specific antibodies and the reliance on recombinant LCMV strains expressing tagged Z proteins (LCMV Z-HA, used in this case) restrict the broader application of Z-based staining for detecting interfering activity. Nonetheless, this observation may offer mechanistic insights into DI-mediated interference, particularly in the context of their ambisense genomic architecture. Mammarenaviruses possess a bi-segmented RNA genome, with each of the S and L segments encoding two genes in opposite orientations (Figure 6). Transcription of the negative-sense open reading frames (ORFs, NP and L) from the genomic RNA occurs first, followed by synthesis of viral complementary antigenomic RNAs that serve as templates for transcription of the positive sense ORFs (GPC and Z). This temporal separation in gene expression is tightly regulated and essential for productive infection [35, 36]. Recent studies have identified deletions within the S segment IGR of arenaviruses associated with high-interference conditions, which correlate with reduced GP expression and impaired viral replication [22, 37]. These deletions may compromise the structural integrity required for transcription termination and reinitiation, thereby silencing the positive-sense genes. This may explain the suppression of Z and GPC translation in cells with high interfering activity, while allowing expression of NP and L from the negative-sense strand [38]. However, there is a lack of empirical evidence linking these genomic findings to viral population phenotypes.

**Figure 6.**
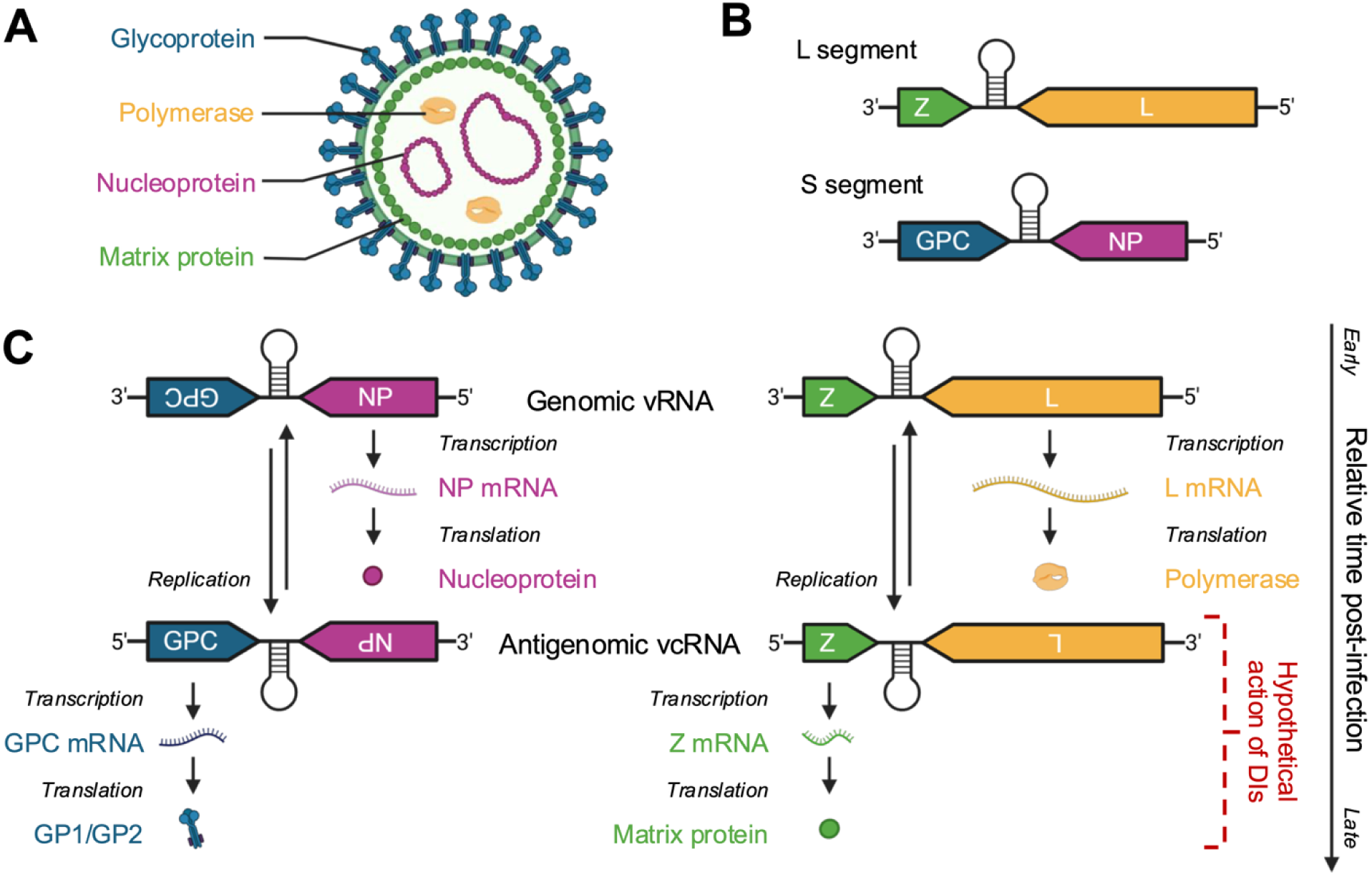
Proposed model of viral interference. (A) Stylized mammarenavirus virion with glycoprotein (GP1/GP2), RNA-dependent RNA polymerase (L), nucleoprotein (NP), and matrix protein (Z) labeled. (B) Genome organization showing ambisense organization for Large (L) segment L and Z genes and Small (S) segment glycoprotein precursor (GPC) and NP genes. (C) Illustration of ambisense replication strategy with hypothesized action of defective interfering particles and/or defective viral genomes on the expression of GP1/GP2 and Z later in infection. Created in BioRender. Honce, R. (2026) https://BioRender.com/rk493qr.

Viral interference, and the associated production of DIPs, has been hypothesized as a virally encoded driver of persistence [4, 23]. There is a cyclical loss and gain of S segment vRNA during persistent infection *in vitro* [35, 39] and less GP expression during persistent infections *in vivo* due to cytoplasmic GP not trafficking to the cell surface [40]. Together, this indicates that the loss of GP expression during high levels of interference mirrors the loss of GP expression that occurs during persistent infections; however, a causal link between these two phenomena has yet to be established [25, 26]. Taken together, these findings support a model in which DI particles and/or DVGs may act at a pre-and/or post-transcriptional but pre-translational stage of the ambisense replication cycle, selectively impairing late gene expression of the positive sense ORFs and halting the production of infectious progeny while still allowing L/NP mediated replication of DVGs. This allowance of early gene expression ensures the minimal replication machinery is present [41] but blocks the expression of GP and Z which halts virion assembly and release, thus creating a non-productive infection carrier state ideal for persistence that may limit host pathology. This temporal disruption may be a key feature of arenavirus biology and could inform future strategies for therapeutic development.

## Methods

### Viruses, cell culture, and antibodies

Lymphocytic choriomeningitis virus (LCMV) Armstrong 53b strain was provided by J. L. Whitton (The Scripps Research Institute) and plasmids for the rescue of recombinant LCMV were provided by Juan Carlos de la Torre (The Scripps Research Institute). Junín virus (JUNV) Candid#1 vaccine strain was provided by R. Tesh (The University of Texas Medical Branch). Recombinant LCMV was generated as previously described [28, 42, 43]. All viruses were propagated in Vero E6 cells which were maintained in Dulbecco’s modified Eagle medium (DMEM) containing 10% fetal bovine serum (FBS), 1% penicillin-streptomycin, and 1% HEPES buffer solution (ThermoFisher). Antibodies for immunostaining were obtained from The Scripps Research Institute Antibody Corse Facility (1-1-3, mouse anti-LCMV nucleoprotein (NP); 33.6, mouse anti-LCMV glycoprotein (GP);) and BEI resources (NA05, mouse anti-JUNV nucleoprotein; BF11, mouse anti-JUNV glycoprotein (GP1)) [44].

### Traditional plaque interfering assay

The traditional plaque interfering assay was completed as previously reported (Figure 1) [27, 28]. Briefly, Vero E6 cells were seeded in a tissue-culture treated 24-well plate at 50,000 cells per well in 500 µL complete DMEM to reach confluency the following day. A sample with unknown interfering activity was placed in a clear microcentrifuge tube (Thermo Scientific, #21-402-904) and exposed to ultraviolet light (254 nM) at 200,000 µJ for 2 minutes to ablate viral replication. Serial dilutions (1:2 to 1:10) of the UV-exposed samples (100 μL) were plated in duplicate and overlaid with 50 PFUs of a known, wild-type LCMV stock (100 μL). After 1.5 hours of adsorption at 37°C, the inoculum was overlaid with 2X DMEM diluted 1:1 with 0.8% agarose in water and returned to the incubator (37°C, 5% CO_2_) for 5 days. After the incubation period, cell monolayers were fixed with the addition of 20% paraformaldehyde in phosphate buffered saline (PBS) for 15 minutes, agarose plugs removed, and plaques counterstained with 1% crystal violet. The enumerated plaques in technical duplicates were averaged for each dilution and plotted against the dilution of unknown sample. The 50% absolute inhibitory concentration (IC_50_) was calculated by a four-parameter logistic curve (GraphPad Prism v10) and standardized to yield interfering activity per milliliter.

### Modified high-throughput focus interference assay

Vero E6 cells were seeded in a white-bottom, 96-well plate at 10,000 cells per well in 100 μL complete DMEM to reach confluency the following day. A sample with unknown interfering activity was placed in a clear microcentrifuge tube (Thermo Scientific, #21-402-904) and exposed to ultraviolet light (254 nM) at 200,000 µJ for 2 minutes to ablate viral replication. Serial dilutions (1:2 to 1:10) of the UV-exposed samples (25 μL) were plated in triplicate and overlaid with 50 FFUs of a known, wild-type LCMV stock (25 μL). After 1.5 hours of adsorption at 37°C, the inoculum was overlaid with 1% carboxymethylcellulose (CMC) in complete DMEM (50 μL) and returned to the incubator (37°C, 5% CO_2_) for 72 hours. After the incubation period, inoculum and CMC overlay were aspirated and monolayers washed twice with 1X PBS, then fixed with the addition of 4% paraformaldehyde in PBS for 15 minutes. Monolayers were then permeabilized with 0.1% triton-X in PBS for 15 minutes (100 μL), washed twice with PBS, then blocked with 5% non-fat dry milk (NFDM) in PBS for 1 hour (100 μL). Viral antigens were detected using an anti-mouse primary antibody specific to the LCMV nucleoprotein (1-1-3; 1:5000 dilution) followed by a peroxidase-conjugated goat anti-mouse secondary antibody (SeraCare #5220-0341; 1:2000 dilution). Viral foci were chromogenized with the addition of a peroxidase substrate (TrueBlue, SeraCare #5510-0030) and quantified on an ImmunoSpot analyzer (CTL, S6 Universal M2). The enumerated foci in technical triplicates were averaged for each dilution and plotted against the dilution of unknown sample. The 50% absolute inhibitory concentration (IC_50_) was calculated by a four-parameter logistic curve (GraphPad Prism v10) and standardized to yield interfering activity per milliliter.

### Interfering activity via glycoprotein immunostaining

For dual determination of focus forming unit (FFU) and focus interfering unit dose-50 (FIU_50_), Vero E6 cells were seeded at 10,000 cells per well in 100 μL complete DMEM in a white bottom 96-well polystyrene plate (FisherScientific #07-200-628) and incubated overnight at 37°C and 5% CO_2_. The following day, serial dilutions (50 μL) of unknown samples were plated in triplicate and allowed to adsorb for 1 hour with gently rocking every 15 minutes to prevent drying of the cellular monolayer. Following adsorption, a 1% CMC matrix overlay diluted in complete DMEM was added to each well (50 μL) and plates were returned to the incubator. After 3 days, the monolayers were fixed using 4% paraformaldehyde in PBS for 15 minutes. Viral antigens were detected using an anti-mouse primary antibody specific to the LCMV glycoprotein (33.6; 1:5000 dilution) followed by a peroxidase-conjugated goat anti-mouse secondary antibody (SeraCare #5220-0341; 1:2000 dilution), followed by the protocol described in the above section for visualization, enumeration of foci, and IC_50_ calculations. The average of enumerated foci in technical triplicates is reported as the log_10_FFU viral titer per milliliter or per gram of sample.

### Quantification of interfering activity by the improved Kärber method

Scanned plate images and/or images binarized in ImageJ [45] were visually assessed for staining patterns. Interference is characterized by lack of GP staining at highest dilutions, followed by foci formation and 1 to 2 countable wells, followed by loss of foci formation due to dilution of sample. Lack of interference is noted by complete staining of cell monolayers at highest dilutions of the unknown sample. Wells positive for interference (i.e., wells above the first countable dilution with distinctly formed foci), were quantified using the Improved Kärber method (https://www.virosin.org/tcid50/TCID50.html).

### Percent maximal staining via imageJ

Use of binarized images and quantification of black pixels in imageJ can improve determination by computational identification of last well(s) positive for interference by identifying the wells with the highest proportion of black pixels. Plates were scanned using the CTL ImmunoSpot analyzer (CTL, S6 Universal M2) or Cytation 7 via Gen5 software (biotek) and images uploaded to imageJ/FIJI [45]. Final plate well images generated using the Cytation 7 were stitched from 4 unique tiled images per well. Using the color threshold tool, the threshold was adjusted so that positively stained foci were segmented as black pixels while background was excluded, and the same minimum and maximum threshold values were used for all wells analyzed on each plate. Thresholding values differed virus-to-virus, plate-to-plate, experiment-to-experiment due to differences in background staining intensity. Following thresholding, images were converted to binary masks (Process > Binary > Make Binary). The percentage of the well area occupied by black pixels was quantified using Analyze > Measure. The percent area of black pixels was recorded as total GP or NP signal per well.

### Quantification of interfering activity via composite score calculation

Foci were quantified on an ImmunoSpot analyzer (CTL, S6 Universal M2). Results from ImmunoSpot also report mean spot sizes, total intensities of spots per well, among other metrics. To calculate DI activity scores for each well or dilution, we used the following formula:

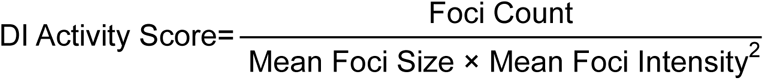

As the ImmunoSpot analyzer does not report mean intensity directly, the formula was modified to:

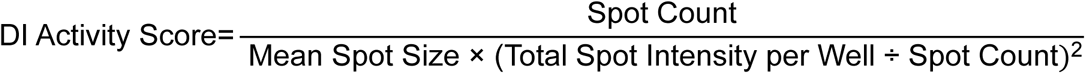

For each well, DI activity scores were plotted against the log_10_ dilution of sample. The 50% absolute inhibitory concentration (IC_50_) was calculated by a four-parameter logistic curve (GraphPad Prism v10) and standardized to yield interfering activity per milliliter.

### Animal infections and sampling

All experimental protocols (PROTO202200022) received approval from the University of Vermont’s Institutional Animal Care and Use Committee (IACUC) and adhered to the guidelines outlined in the Guide for the Care and Use of Laboratory Animals. C57BL6 mice (Jackson) were maintained under a 12-hour light/dark cycle at a stable ambient temperature of 27°C and controlled humidity. Throughout the experiment, they had unrestricted access to food and water and were co-housed (2-4 per cage based on like infection status and sex) in isolation units equipped with HEPA filtration maintained under ventilated negative pressure. Persistent infections were established by intraperitoneal injection of neonatal mice within the first 24 hours of birth with 1000 plaque-forming units (PFUs) of recombinant LCMV Armstrong 53b suspended in phosphate buffered saline (PBS) [46]. Sham, uninfected control animals were injected with PBS only. No randomization was used to assign litters to infection or control. For samples reported in this study, serum was collected at week 12 of life via terminal cardiac puncture. Samples were left to clot at room temperature for 30 minutes, followed by centrifugation at 2,000 × g for 15 minutes at 4°C to separate the serum. The resulting serum was assigned a unique experimental code for sample blinding and stored at −80°C. All procedures took place within a biosafety level 3 (BSL-3) facility. Researchers were required to wear Versaflo PAPR respiratory protection (3M) with all mouse handling and tissue culture performed exclusively inside Class 2A biosafety cabinets. Data displayed is representative of a larger mouse colony maintained across two independent 10-month periods across 2 years encompassing n=90 persistently infected and n=46 sham infected male and female mice used for other experiments. Inclusion of mice in this study were based on adequate volume of serum available for testing across the assays reported herein and previous positive infectious titer determination.

## Notes

### Competing Interest Statement

The authors have declared no competing interest.

### Funding

- National Institute of Allergy and Infectious Diseases: R01AI171408
- National Heart, Lung, and Blood Institute: T32HL076122

## References

1. Magnus PV. Studies on Interferenee in Experimental Influenza. I. Biological Observations. *Arkiv for Kemi*, Mineralogi och Geologi 1947;24B(7):6.

2. von MP. Propagation of the PR8 strain of influenza A virus in chick embryos. II. The formation of incomplete virus following inoculation of large doses of seed virus. Acta Pathol Microbiol Scand 1951;28(3):278–293.

3. Cooper PD, Bellett AJ. A transmissible interfering component of vesicular stomatitis virus preparations. J Gen Microbiol 1959;21:485–497.

4. Huang AS. Defective interfering viruses. Annu Rev Microbiol 1973;27:101–117.

5. Lopez CB. Defective viral genomes: critical danger signals of viral infections. J Virol 2014;88(16):8720–8723.

6. Sun Y, Jain D, Koziol-White CJ, Genoyer E, Gilbert M et al. Immunostimulatory Defective Viral Genomes from Respiratory Syncytial Virus Promote a Strong Innate Antiviral Response during Infection in Mice and Humans. PLoS Pathog 2015;11(9):e1005122.

7. Vasilijevic J, Zamarreno N, Oliveros JC, Rodriguez-Frandsen A, Gomez G, et al. Reduced accumulation of defective viral genomes contributes to severe outcome in influenza virus infected patients. PLoS Pathog 2017;13(10):e1006650.

8. Pesko KN, Fitzpatrick KA, Ryan EM, Shi PY, Zhang B, et al. Internally deleted WNV genomes isolated from exotic birds in New Mexico: function in cells, mosquitoes, and mice. Virology 2012;427(1):10–17.

9. Yount JS, Kraus TA, Horvath CM, Moran TM, Lopez CB. A novel role for viral-defective interfering particles in enhancing dendritic cell maturation. J Immunol 2006;177(7):4503–4513.

10. Kalamvoki M, Norris V. A Defective Viral Particle Approach to COVID-19. Cells 2022;11(2):302.

11. Pitchai FNN, Tanner EJ, Khetan N, Vasen G, Levrel C, et al. Engineered deletions of HIV replicate conditionally to reduce disease in nonhuman primates. Science 2024;385(6709):eadn5866.

12. Rezelj VV, Carrau L, Merwaiss F, Levi LI, Erazo D, et al. Defective viral genomes as therapeutic interfering particles against flavivirus infection in mammalian and mosquito hosts. Nat Commun 2021;12(1):2290.

13. Welch SR, Spengler JR, Harmon JR, Coleman-McCray JD, Scholte FEM, et al. Defective Interfering Viral Particle Treatment Reduces Clinical Signs and Protects Hamsters from Lethal Nipah Virus Disease. mBio 2022;13(2):e0329421.

14. Alnaji FG, Holmes JR, Rendon G, Vera JC, Fields CJ, et al. Sequencing Framework for the Sensitive Detection and Precise Mapping of Defective Interfering Particle-Associated Deletions across Influenza A and B Viruses. J Virol 2019;93(11).

15. Beauclair G, Mura M, Combredet C, Tangy F, Jouvenet N, et al. DI-tector: defective interfering viral genomes’ detector for next-generation sequencing data. RNA 2018;24(10):1285–1296.

16. Bosma TJ, Karagiannis K, Santana-Quintero L, Ilyushina N, Zagorodnyaya T, et al. Identification and quantification of defective virus genomes in high throughput sequencing data using DVG-profiler, a novel post-sequence alignment processing algorithm. PLoS One 2019;14(5):e0216944.

17. Boussier J, Munier S, Achouri E, Meyer B, Crescenzo-Chaigne B, et al. RNA-seq accuracy and reproducibility for the mapping and quantification of influenza defective viral genomes. RNA 2020;26(12):1905–1918.

18. Jaworski E, Routh A. Parallel ClickSeq and Nanopore sequencing elucidates the rapid evolution of defective-interfering RNAs in Flock House virus. PLoS Pathog 2017;13(5):e1006365.

19. Lui WY, Yuen CK, Li C, Wong WM, Lui PY, et al. SMRT sequencing revealed the diversity and characteristics of defective interfering RNAs in influenza A (H7N9) virus infection. Emerg Microbes Infect 2019;8(1):662–674.

20. Wang C, Forst CV, Chou TW, Geber A, Wang M, et al. Cell-to-Cell Variation in Defective Virus Expression and Effects on Host Responses during Influenza Virus Infection. mBio 2020;11(1).

21. Wang C, Honce R, Salvatore M, Chow D, Randazzo D, et al. Influenza Defective Interfering Virus Promotes Multiciliated Cell Differentiation and Reduces the Inflammatory Response in Mice. J Virol 2023;97(6):e0049323.

22. Hackbart M, Lopez CB. Characterization of non-standard viral genomes during arenavirus infections identifies prominent S RNA intergenic region deletions. mBio 2024;15(10):e0161224.

23. Meyer BJ, Southern PJ. A novel type of defective viral genome suggests a unique strategy to establish and maintain persistent lymphocytic choriomeningitis virus infections. J Virol 1997;71(9):6757–6764.

24. Southern PJ, Dyrberg T, Schwimmbeck PL, Oldstone MB. Persistent virus infection and development of virus-induced disease. J Autoimmun 1990;3 Suppl 1:13–20.

25. Francis SJ, Southern PJ. Deleted viral RNAs and lymphocytic choriomeningitis virus persistence in vitro. J Gen Virol 1988;69 (Pt 8)(8):1893–1902.

26. Francis SJ, Southern PJ. Molecular analysis of viral RNAs in mice persistently infected with lymphocytic choriomeningitis virus. J Virol 1988;62(4):1251–1257.

27. Ziegler CM, Eisenhauer P, Bruce EA, Beganovic V, King BR, et al. A novel phosphoserine motif in the LCMV matrix protein Z regulates the release of infectious virus and defective interfering particles. J Gen Virol 2016;97(9):2084–2089.

28. Ziegler CM, Eisenhauer P, Bruce EA, Weir ME, King BR, et al. The Lymphocytic Choriomeningitis Virus Matrix Protein PPXY Late Domain Drives the Production of Defective Interfering Particles. PLoS Pathog 2016;12(3):e1005501.

29. Popescu M, Schaefer H, Lehmann-Grube F. Homologous interference of lymphocytic choriomeningitis virus: detection and measurement of interference focus-forming units. J Virol 1976;20(1):1–8.

30. Iwasaki M, Ngo N, Cubitt B, de la Torre JC. Efficient Interaction between Arenavirus Nucleoprotein (NP) and RNA-Dependent RNA Polymerase (L) Is Mediated by the Virus Nucleocapsid (NP-RNA) Template. J Virol 2015;89(10):5734–5738.

31. Morin B, Coutard B, Lelke M, Ferron F, Kerber R, et al. The N-Terminal Domain of the Arenavirus L Protein Is an RNA Endonuclease Essential in mRNA Transcription. PLOS Pathogens 2010;6(9):e1001038.

32. King BR, Hershkowitz D, Eisenhauer PL, Weir ME, Ziegler CM, et al. A Map of the Arenavirus Nucleoprotein-Host Protein Interactome Reveals that Junin Virus Selectively Impairs the Antiviral Activity of Double-Stranded RNA-Activated Protein Kinase (PKR). J Virol 2017;91(15):e00763–00717.

33. Gowen BB, Hickerson BT, York J, Westover JB, Sefing EJ, et al. Second-Generation Live-Attenuated Candid#1 Vaccine Virus Resists Reversion and Protects against Lethal Junin Virus Infection in Guinea Pigs. J Virol 2021;95(14):e0039721.

34. Palacios G, Druce J, Du L, Tran T, Birch C, et al. A new arenavirus in a cluster of fatal transplant-associated diseases. N Engl J Med 2008;358(10):991–998.

35. King BR, Samacoits A, Eisenhauer PL, Ziegler CM, Bruce EA et al. Visualization of Arenavirus RNA Species in Individual Cells by Single-Molecule Fluorescence In Situ Hybridization Suggests a Model of Cyclical Infection and Clearance during Persistence. J Virol 2018;92(12):e02241–02217.

36. Kang H, Cong J, Wang C, Ji W, Xin Y, et al. Structural basis for recognition and regulation of arenavirus polymerase L by Z protein. Nat Commun 2021;12(1):4134.

37. Gass JT, Nofchissey RA, Twohig FM, Ye C, Goodfellow SM, et al. Discovery of a novel lymphocytic choriomeningitis virus strain associated with severe human disease in immunocompetent patient, New Mexico. Emerg Microbes Infect 2025;14(1):2542250.

38. Labudova M, Tomaskova J, Skultety L, Pastorek J, Pastorekova S. The nucleoprotein of lymphocytic choriomeningitis virus facilitates spread of persistent infection through stabilization of the keratin network. J Virol 2009;83(16):7842–7849.

39. Welsh RM, Jr., Buchmeier MJ. Protein analysis of defective interfering lymphocytic choriomeningitis virus and persistently infected cells. Virology 1979;96(2):503–515.

40. Oldstone MB, Buchmeier MJ. Restricted expression of viral glycoprotein in cells of persistently infected mice. Nature 1982;300(5890):360–362.

41. Lee KJ, Novella IS, Teng MN, Oldstone MB, de La Torre JC. NP and L proteins of lymphocytic choriomeningitis virus (LCMV) are sufficient for efficient transcription and replication of LCMV genomic RNA analogs. J Virol 2000;74(8):3470–3477.

42. Flatz L, Bergthaler A, de la Torre JC, Pinschewer DD. Recovery of an arenavirus entirely from RNA polymerase I/II-driven cDNA. Proc Natl Acad Sci U S A 2006;103(12):4663–4668.

43. Ziegler CM, Bruce EA, Kelly JA, King BR, Botten JW. The use of novel epitope-tagged arenaviruses reveals that Rab5c-positive endosomal membranes are targeted by the LCMV matrix protein. J Gen Virol 2018;99(2):187–193.

44. Sanchez A, Pifat DY, Kenyon RH, Peters CJ, McCormick JB, et al. Junin virus monoclonal antibodies: characterization and cross-reactivity with other arenaviruses. J Gen Virol 1989;70 (Pt 5):1125–1132.

45. Stossi F, Singh PK. Basic Image Analysis and Manipulation in ImageJ/Fiji. Curr Protoc 2023;3(7):e849.

46. von Herrath M, Whitton JL. Animal models using lymphocytic choriomeningitis virus. Curr Protoc Immunol 2001;Chapter 19:Unit 19 10.

